# Microsaccades are directed towards the midpoint between targets in a variably cued attention task

**DOI:** 10.1101/2023.01.24.525403

**Authors:** Shawn M. Willett, J. Patrick Mayo

## Abstract

Reliable, non-invasive biomarkers that reveal the internal state of a subject are an invaluable tool for neurological diagnoses. Small fixational eye movements, called microsaccades, are a candidate biomarker thought to reflect a subject’s focus of attention (1, 2). The linkage between the direction of microsaccades and attention has mainly been demonstrated using explicit and unambiguous attentional cues. However, the natural world is seldom predictable and rarely provides unambiguous information. Thus, a useful biomarker must be robust to such changes in environmental statistics. To determine how well microsaccades reveal visual-spatial attention across behavioral contexts, we analyzed these fixational eye movements in monkeys performing a conventional change detection task. The task included two stimulus locations and variable cue validities across blocks of trials. Subjects were adept at the task, showing precise and graded modulations of visual attention for subtle target changes and performing better and faster when the cue was more reliable (3). However, over tens of thousands of microsaccades, we found no difference in microsaccade direction between cued locations when cue variability was high nor between hit and miss trials. Instead, microsaccades were made towards the midpoint of the two target locations, not towards individual targets. Our results suggest that the direction of microsaccades should be interpreted with caution and may not be a reliable measure of covert spatial attention in more complex viewing conditions.

**Significance Statement:** Small fixational eye movements called microsaccades are thought to “point” towards a location that is being attended in the visual periphery. This phenomenon has largely been studied using visual cues that unambiguously indicate the location of the upcoming stimulus change. Because the natural world is rarely unambiguous, we studied the relationship between microsaccade direction and spatial attention using less reliable cues. We found that monkeys’ microsaccade directions in a standard visuospatial attention task did not indicate the animals’ focus of attention, despite behavioral and neuronal evidence of spatial attention. Instead, microsaccades were made towards the midpoint between the target locations in both animals, suggesting a more complex relationship between microsaccades and attention in naturalistic settings.

## Introduction

The noninvasive diagnosis of disease states is a central goal of translational science. The ability to measure small changes in physiology as an alternative to expensive and invasive metrics is essential to improving the affordability, accessibility, and efficiency of treatment. Given the critical role that foveated viewing plays in primate vision and the ability to measure eye position with high precision, it is not surprising that eye movements have been researched as potential biomarkers for neurological disease and cognitive states (4, 5), among other alternatives (6). Because humans spend roughly 80% of their time fixating (7), fixational eye movements such as “microsaccades” are of particular interest as a candidate biomarker.

Microsaccades are small, rapid eye movements that occur during fixation. They contribute to multiple aspects of visual function including correcting for small errors in eye position (8), refreshing the activity of the visual system (9-11), enhancing fine spatial acuity (12, 13, though see 14), and interacting with the allocation of visual-spatial attention (for review see 15). Indeed, microsaccade rate and amplitude have been proposed as a biomarkers for many neurological diseases including autism spectrum disorder, schizophrenia, and attention-deficit hyperactivity disorder (for review see 16). Microsaccades have also been linked to phenomena that are more subtle than neurological impairments such as covert visual-spatial attention.

There is an extensive body of research linking microsaccades to spatial attention (1, 2, 17-21). Early work by Hafed and Clark (1) showed that microsaccades are directed towards the direction of a preceding attentional cue in a covert attention task. Thus, the microsaccades appeared to “point” towards the cued location and the presumed focus of attention. This result and supporting work (2, 17, 19, 22) suggest that microsaccades could be used to reveal the otherwise hidden locus of visuo-spatial attention while the eyes remain relatively still.

Motivated by the finding that the direction of microsaccades aligns with cued locations in deterministic tasks, we set out to investigate this relationship in scenarios more comparable to everyday experiences. Natural environments provide an abundance of cues that signal uncertain future events. For example, the brief appearance and disappearance of an animal from behind a tree indicates some likelihood that the animal will appear again at the same location. But it would be foolish to assume that the animal’s re-appearance is entirely predictable in space and time. Instead, we must integrate and maintain knowledge of the world to adaptively deploy attention. In fact, previous work has shown that microsaccade direction can be biased by cues stored in memory (23). But little is known about how variable cue validity affects microsaccade direction (though see 2). The usefulness of microsaccades as a biomarker for covert spatial attention requires a more systematic understanding of the interaction of microsaccade direction and attention under more variable experimental conditions.

Here we sought to determine how variable cueing affected the interaction of visual-spatial attention and the direction of microsaccades. We measured microsaccades while monkeys fixated during an orientation change detection task with a target in each visual hemifield (3). We found that microsaccades were directed towards the midpoint between the two targets. There was a slight bias towards the cued location in the less variable cueing regime and this bias was not present in the more variable cueing regime. Importantly, there was no difference in microsaccade direction for correctly detected versus missed trials despite considerable changes in performance and reaction times (i.e., canonical behavioral correlates of visual attention). Our results suggest that the usefulness of microsaccades as a biomarker of visual attention may be limited in naturalistic scenarios that contain less predictable events.

## Results

To investigate the relationship between microsaccades and visual spatial attention, we trained monkeys to perform an orientation change detection task. The task required monkeys to report a change in the orientation of one of two simultaneously presented Gabor patches (“targets”) after a random fixation time (Fig. 1A; see Methods). The target locations and orientations before the change remained constant in each session. One target appeared in the left visual field and the other appeared in the right visual field. Target locations varied across sessions (Fig. 1B), but were restricted to the lower visual field because of accompanying neurophysiological recordings in cortical area V4 (3). Behavioral sessions consisted of 50-trial blocks, alternating between each of the cuing conditions. Each block was preceded by a small number of instruction trials where a visual cue was presented at the target location where the orientation change was likely to occur in the immediately following block. After successfully completing the instruction trials, the visual cue was removed for the rest of the block and monkeys were required to maintain the cued location in memory. Eye movements were recorded using scleral search coils (24), and instruction trials were omitted from analyses.

**Figure 1.**
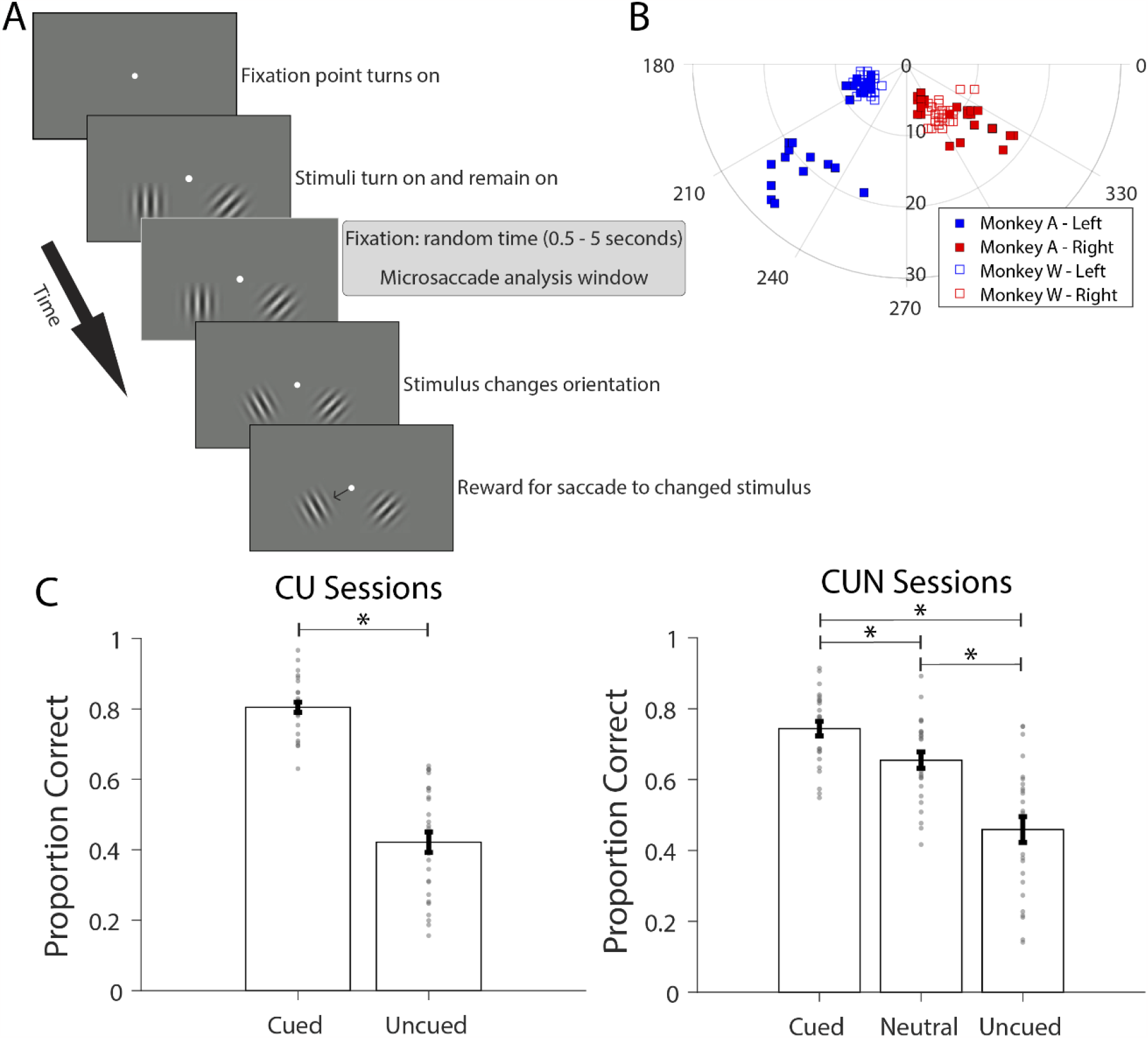
Behavioral task, target locations and performance. A) Schematic showing the progression of a single example trial. Microsaccades were analyzed during the sustained fixation period (shaded text box) while monkeys foveated a central fixation point (white dot) before making a saccade (thin black arrow) to the stimulus that changed orientation (bottom). B) Target locations for monkey A (filled squares) and monkey W (open squares). The angular values correspond to the angular position of the target relative to fixation (0°,0°), while the radial values correspond to the eccentricity of the target (°). C) The proportion of correct trials for Cued/Uncued (CU; left panel) and Cued/Uncued/Neutral (CUN; right panel) sessions across monkeys. The height of each bar and error bars correspond to the mean ± SEM performance across sessions (CU n = 29; CUN n = 26). Small grey dots represent performance for each session. Asterisks indicate that Cued performance is significantly higher than Uncued performance in both CU and CUN cueing regimes, and Neutral Cued performance is significantly higher than Uncued performance in the CUN regime (two-tailed paired t-test, Dunn-Bonferroni corrected, *p* < 0.0125). CU sessions had on average 987 ± 47 SEM trials per session; CUN sessions had on average 1040 ± 30 trials per session.

Each session utilized one of two possible cueing regimes: Cued/Uncued (referred to as ‘CU’) or Cued/Uncued/Neutral (referred to as ‘CUN’). The cued and uncued target locations alternated within a session in 50-trial blocks. During Cued/Uncued blocks, the target at the cued location had an 80% likelihood of changing its orientation while the target at the uncued location had a 20% likelihood. Therefore, the CU regime only consisted of Cued/Uncued blocks with 80% cueing validity. During Neutral blocks, each target had a 50% likelihood of changing its orientation (i.e., cue presented at both stimulus locations during instruction trials). Therefore, the CUN regime consisted of both Cued/Uncued blocks and Neutral blocks with 80/50% cue validities. Sessions consisted of either CU or CUN cueing regimes.

Monkeys performed the task well and displayed behavioral benefits of attention in the form of increased behavioral performance (Fig. 1C). In prior work, we found lower discrimination thresholds, and faster reaction times at the cued location in the CUN regime (3). In the 26 sessions using the CUN regime, monkeys’ proportion correct was higher on Cued trials relative to Neutral and Uncued trials (Cued mean = 0.74 ± SEM = 0.02 vs. Neutral mean = 0.66 ± SEM = 0.02; *p* = 5.41 × 10^−8^; Cued vs. Uncued mean = 0.46 ± SEM = 0.04; *p* = 5.41 × 10^−8^; Dunn-Bonferroni multiple comparison threshold, p = 0.0125) and Neutral trials relative to Uncued trials (Neutral mean = 0.66 ± SEM = 0.02 vs. Uncued mean = 0.46 ± SEM = 0.04; *p* = 2.05 × 10^−11^). A similar effect was found in the 29 sessions using the CU regime (Cued mean = 0.81 ± SEM = 0.01 vs. Uncued mean = 0.42 ± SEM = 0.03; *p* = 6.37 × 10^−13^; Dunn-Bonferroni multiple comparison threshold, p = 0.0125). Overall, these findings indicate that monkeys exhibited significant attention-related improvements in behavioral performance as a function of the cue validity.

We measured microsaccades during the fixation period between stimulus onset and target change while the monkeys attended to one of the targets to detect the orientation change (Fig. 1A). We classified eye movements as microsaccades if the movement amplitude was less than 2° and the velocity surpassed 6° per second (see Methods). Our results were robust to changes in velocity threshold (5-8 °/s) and amplitude threshold (1° or 0.5°). We first verified that microsaccades detected in our task displayed three commonly found characteristics (2, 25). First, microsaccades in both monkeys fell along the main sequence (Fig. 2A & D), indicating that microsaccades were truly smaller amplitude versions of saccades and the relationship between peak movement amplitude and peak movement velocity (“main sequence”) was maintained across saccade types (26). Second, microsaccades were typically a fraction of a degree in amplitude (Fig. 2B & E) (2, 25, 27). Finally, stimulus onset led to a stereotypical suppression and rebound in the number of microsaccades, also known as the microsaccade rate signature (2, 25, 28, 29) (Fig. 2C & F). Thus, our methods for measuring microsaccade dynamics agreed with well-established microsaccade characteristics.

**Figure 2.**
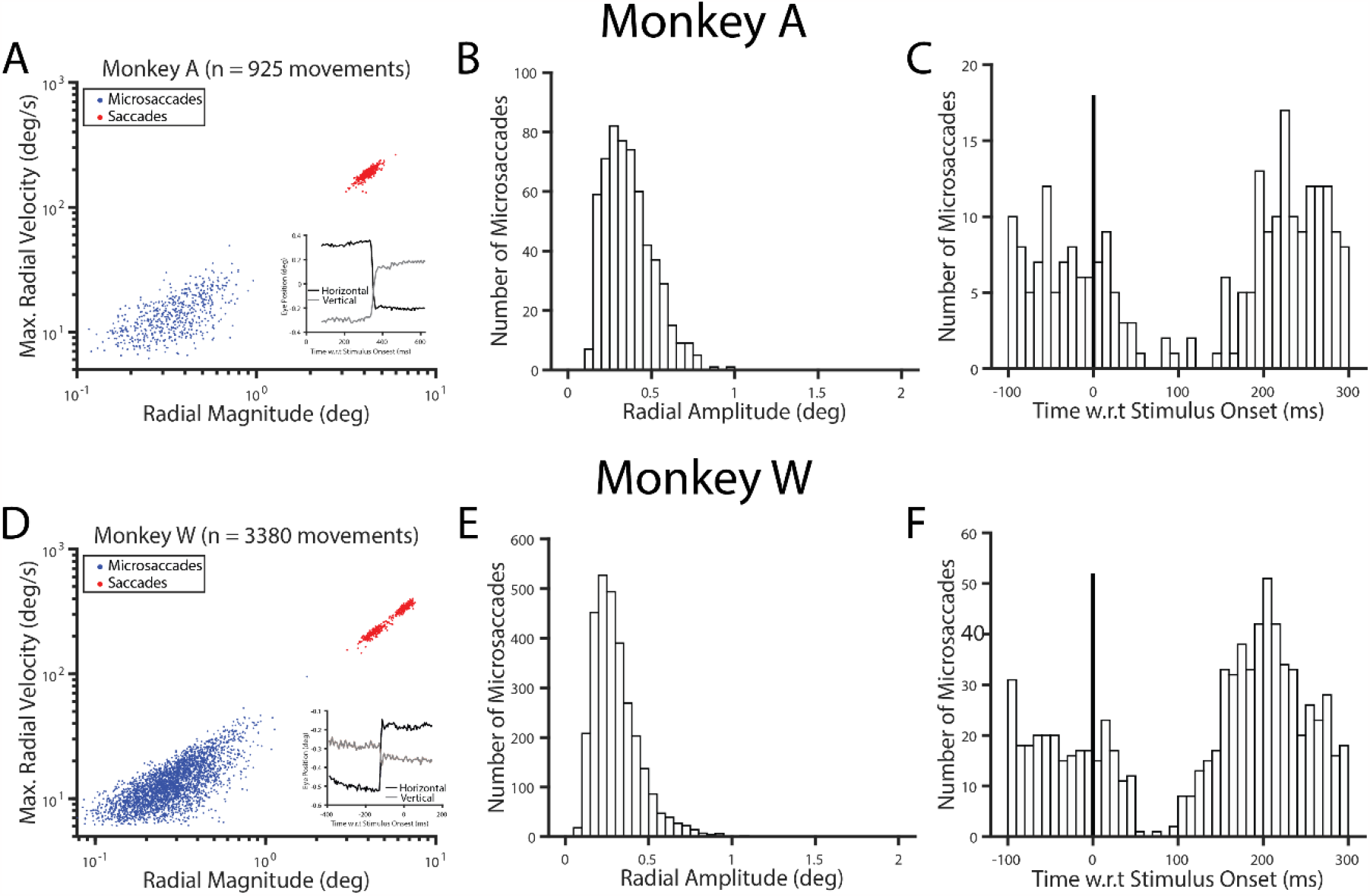
Microsaccades fall along the main sequence and exhibit typical amplitudes and post-stimulus dynamics. A) The main sequence plot for monkey A from a representative session, including all microsaccades after stimulus onset. Red points indicate saccades (movements ≥ 2°) and blue points indicate microsaccades (movements < 2°). The inset is an example microsaccade from monkey A directed up and to the left. B) The distribution of microsaccade amplitudes after stimulus onset but prior to target change for the same representative session in A. C) The number of microsaccades aligned to stimulus onset for the same representative session in A (bin width = 10 ms). This plot includes microsaccades prior to stimulus onset (< 0 ms) and shows the classic microsaccade rate suppression effect (2, 25, 28, 29). D-F) Same as A-C but for a representative session from monkey W.

In tasks that present a 100% valid cue, microsaccades tend to “point” to the cued target location. That is, the peak of the distribution of microsaccade directions is nonuniform and strongly biased toward the attended target (1, 20, 23). We performed similar analyses on our data using variable cueing (<100%). Figure 3A shows the distribution of microsaccade directions for each monkey in an example session using the CUN regime. In each panel, angular position around the circle corresponds to the angular direction of the microsaccade, and the amplitude of each bin corresponds to the number of microsaccades made in that direction. Qualitatively, these microsaccade direction distributions display two major features. First, as expected, the distributions were largely bimodal because microsaccades are often made away from and then back towards fixation (1, 20). Second, in both monkeys microsaccades were directed not towards the target locations themselves, but towards the midpoint between the two targets (Fig. 3A; red and blue squares). This finding appeared to be independent of the cueing condition.

**Figure 3.**
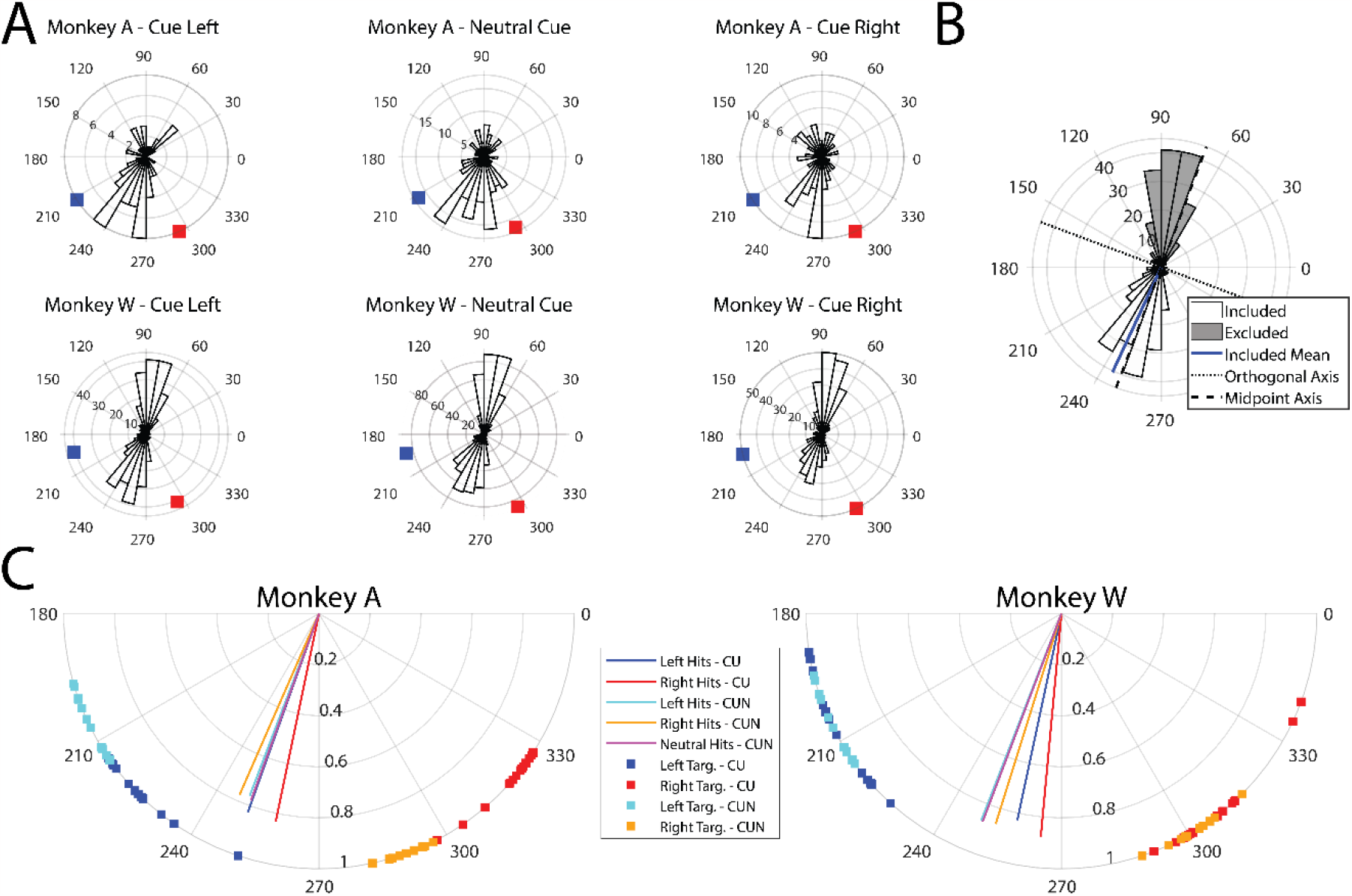
Microsaccades are directed between target locations. A) Example microsaccade direction distributions to different cueing conditions (left to right) from a single CUN session for monkey A (top row) and for monkey W (bottom row). The blue rectangle corresponds to the left stimulus’ angular position while the red rectangle corresponds to the right stimulus’ angular position. The bin width is 10° and bin height (radius) is the number of microsaccades that occur in that direction. B) A schematic showing the included direction distribution (white bins) used in subsequent analyses, and the solid blue line indicating the circular mean of the included distribution. We found the midpoint between both targets (along the dashed line) and took all microsaccades that fell below the orthogonal axis (dotted line). C) The average microsaccade direction across sessions for monkey A (left panel) and monkey W (right panel). Data are separated by cueing regime (Monkey A: n=14/14 CN/CUN sessions; Monkey W: n=15/12 CN/CUN sessions) and cued location (left vs. right vs. neutral). The angle of each vector corresponds to the angular direction of microsaccades averaged across sessions. The length of each vector signifies the variance of the distribution with lengths of one indicating no variance and lengths of zero corresponding to uniformly distributed data around the circle. Squares along outer ring correspond to the angular position of the targets across sessions. Note that the legend in panel C does not apply to panels A and B to keep panels A and B consistent with coloring in Figure 1B and 4A.

To summarize microsaccade directions across sessions, we focused on movements in the hemifield containing the targets because movements in the opposite direction are thought to reacquire the fixation point (due to ocular drift or earlier microsaccades toward the targets). As illustrated in Figure 3B, this separation was achieved by drawing a line along the Euclidean midpoint between the target locations in each session, and then constructing an axis orthogonal to the midline. We retained microsaccades that fell below this axis orthogonal to the midpoint between the two targets (Fig. 3B). Results were robust if we used an angular midpoint. The circular mean for each condition of each session was calculated using the Circular Statistics toolbox (30). Figure 3C shows the average resultant vector across sessions for each condition and monkey (Monkey A: n=14/14 CN/CUN sessions; Monkey W: n=15/12 CN/CUN sessions). The angle of the vector indicates the average microsaccade direction, and the length of the vector summarizes the variance of microsaccade direction (Variance = 1 – Resultant Vector Length). Lengths of 1 indicate that all session means lie along the same angle while lengths of 0 indicate a uniform distribution of microsaccade direction. Consistent with the representative sessions in Figure 3A, averaged microsaccade directions across sessions point toward the midpoint of the two target distributions in both cueing regimes (CU and CUN), regardless of condition. Thus, despite pronounced cue-related improvements in behavior across sessions (Fig. 1C), these results reveal that both monkeys directed their microsaccades towards a location between the two targets.

Because target locations varied somewhat across sessions, we wanted to ensure that the microsaccades were directed towards the true midpoint between the two targets. The average microsaccade direction across sessions above (Fig. 3C) does not quantify the proximity of the session-by-session average to the midpoint. We therefore normalized the microsaccade directions so that they fell along an equidistant scale centered between the targets of each session (Fig 4A). To do this, we developed a metric to quantify how microsaccade direction changed as a function of target positions. We normalized the distance to the midpoint between the targets (see Methods) so that values of -1 indicate microsaccade directions aligned to the left target, values of 1 indicate microsaccade directions aligned to the right target, and values of 0 indicate microsaccade directions aligned to the Euclidean midpoint between the two targets. Due to the similarity in behavior across both monkeys (Fig. 3C) and because this metric accounted for the differences in target positions across sessions, we combined our data across monkeys.

**Figure 4.**
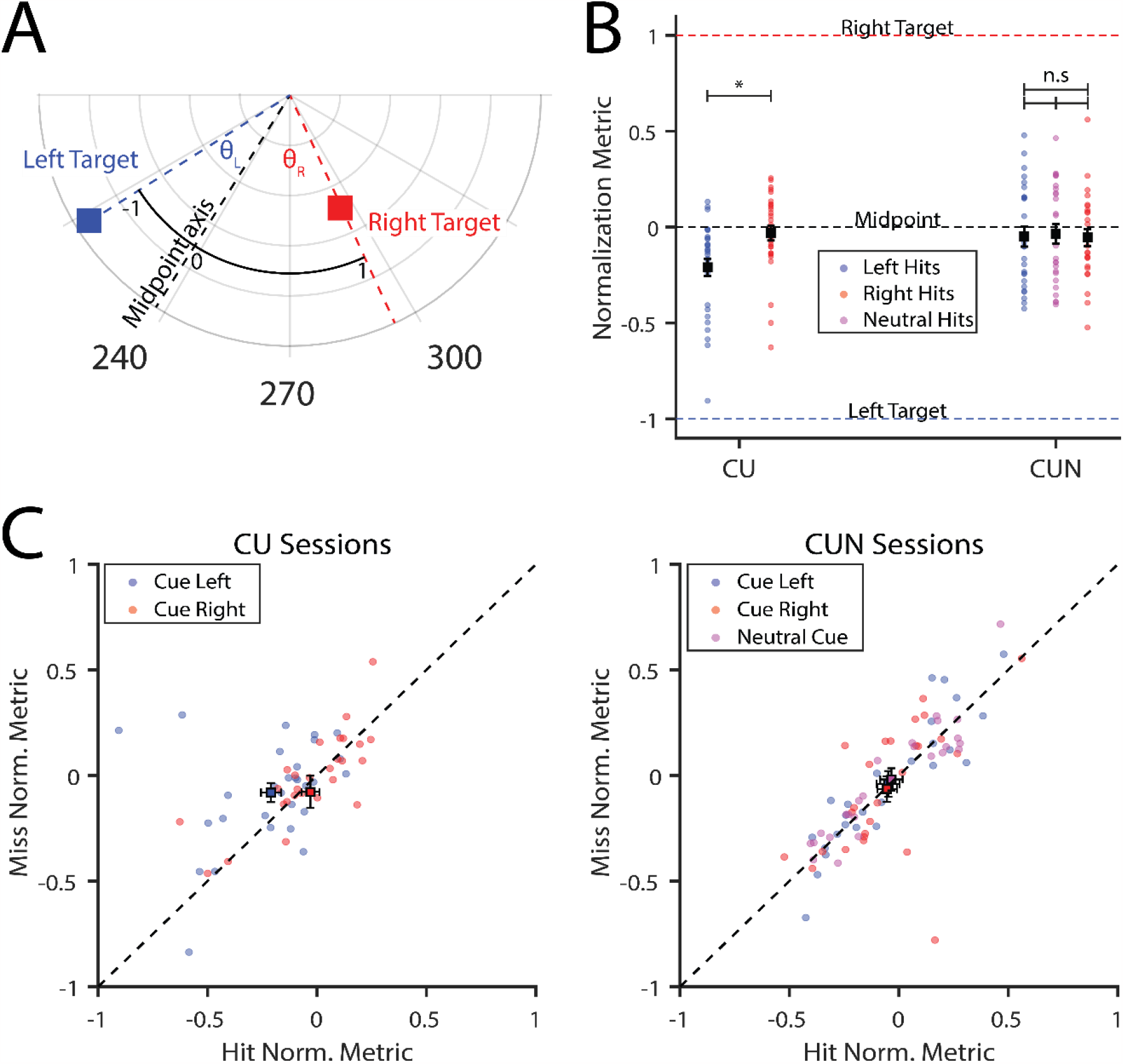
Microsaccades are directed to the midpoint, display less bias in the more variable cue regime, and do not differentiate hit and miss trials. A) A schematic showing the interpretation of the normalization metric. B) The normalization metric across monkeys for different cued directions (left vs. right vs. neutral) for CU (n = 29 sessions) and CUN (n = 26 sessions) cueing regimes. Black squares with error bars represent mean ± SEM. Data points are the average normalization metric values per session. Right hits are significantly more positive than left hits only for the CU sessions and there are no significant differences across cued conditions in CUN sessions. Asterisks indicate significant differences across conditions (two-tailed paired t-test, Dunn-Bonferroni corrected, *p* < 0.0125; n.s indicates no significance). Dashed horizontal lines reference key values for interpretating the normalization metric (see Methods). C) Comparison between the normalization metric values for the average hit and miss trials per session for different cueing regimes (CU, left panel; CUN, right panel). The dashed line indicates the unity line. One data point was removed for visualization but included in all statistical tests (CU panel, hit = 0.037 and miss =-1.855). Note for CU sessions there were no right miss trials in one session so n = 28/29 sessions. There was no significant difference between the hit and miss distributions for either cueing regime.

Figure 4B shows that normalized microsaccade directions were concentrated around zero and aligned to the Euclidean midpoint between the two targets. But are microsaccades biased by the cued location around this midpoint? In the CU sessions, we found that normalized directions were significantly more negative (i.e., towards the left target) for correctly reported cued-left targets and more positive (i.e., towards the right target) for correctly reported cued-right targets (CU Cued Left mean = -0.21 ± SEM = 0.05 vs. CU Cued Right mean = 0.03 ± SEM = 0.04; *p* = 9.56 × 10^−5^; Dunn-Bonferroni multiple comparison threshold, p = 0.0125). The slight change in microsaccade direction in the CU cueing regime corresponds to a shift of roughly 10% towards the cued location relative to the average microsaccade direction of the session.

In the CUN sessions where cue validity was more variable, there was no significant bias in microsaccade direction. Figure 4B shows that microsaccade directions for trials with correctly reported left or right targets did not differ from one another or from the Neutral trials (CUN Cued Left mean = -0.05 ± SEM = 0.05 vs. CUN Cued Right mean = -0.05 ± SEM = 0.05; *p* = 0.87; CUN Cued Left vs. CUN Neutral Cue mean = -0.03 ± SEM = 0.05; *p* = 0.57; CUN Cued Right vs. CUN Neutral Cue; *p* = 0.60; ; Dunn-Bonferroni multiple comparison threshold, p = 0.0125). Thus, the CUN sessions that contained two different cue validities (80/50%) resulted in consistently unbiased microsaccade directions across the conditions. These results suggest that as cue validities become more variable across blocks microsaccade directions are less biased towards the cued location, even in blocks with the same cue validity (Cued/Uncued blocks on CU session vs Cued/Uncued blocks on CUN sessions).

If microsaccades are an indicator of the locus of covert attention (1, 2), then the microsaccade direction should be more aligned with the cued target for trials in which monkeys correctly detected orientation change (“hits”) versus trials in which monkeys missed the change (“misses”). To directly test this notion, we plotted the distribution of normalized microsaccade directions for hits against misses (Fig. 4C). Across cueing regimes, microsaccade direction did not significantly differ between trials in which monkeys correctly detect vs. miss the orientation change (CU Left Hit vs. CU Left Miss mean = -0.08 ± SEM 0.05; *p* = 0.03; CU Right Hit vs. CU Right Miss mean = -0.08 ± SEM 0.08; *p* = 0.52; CUN Left Hit vs. CUN Left Miss mean = -0.04 ± SEM = 0.06; *p* = 0.74; CUN Right Hit vs. CUN Right Miss mean = -0.06 ± SEM 0.06; *p* = 0.84; CUN Neutral Hit vs. CUN Neutral Miss mean = -0.02 ± SEM = 0.05; *p* = 0.43; ; Dunn-Bonferroni multiple comparison threshold, p = 0.01). Interestingly, the clear relationship between hit and miss direction (data falling near the unity line) may suggest that the neural population generating microsaccades is unaffected by attention (see Discussion). Thus, we failed to find evidence supporting the idea that microsaccade direction reflects the focus of attention.

We next performed a series of control analyses to rule out alternative explanations of our data. First, we investigated whether microsaccades were biased toward the vertical meridian in the lower hemifield, as opposed to the midpoint between targets. This behavior was a possibility since targets were always presented below the horizontal azimuth. To test this alternative, we measured the number of times in which the midpoint or vertical meridian fell within the 95% confidence intervals (95% C.I.) of the microsaccade distribution for a given session-condition combination. Of the 136 conditions (29 CU sessions x 2 condition + 26 CUN sessions x 3 conditions), 100% of the microsaccade distributions (n = 136/136) contained the session midpoint within the 95% C.I. whereas only ∼14% of the microsaccade distributions (n = 19/136) contained the vertical meridian within the 95% C.I. Approximately 63% (n = 12/19) of the microsaccade distributions that contained the vertical meridian were within 15° of the midpoint between the two targets. These results confirm that the monkeys were making microsaccades to the midpoint and not simply making purely downward microsaccades to along the vertical meridian.

Second, we asked if the absence of microsaccades aligned to the target locations could be due to a bias in fixation position (31). For example, subjects may have been less likely to make microsaccades towards a particular target location if their fixation position was already biased in that direction. To better understand the relationship between fixation and microsaccade direction, we looked at the average fixation position during the period between stimulus onset and prior to target change (Fig. 1A, gray shading). Across both monkeys, cueing condition did not lead to a substantial difference in fixation position (Fig. 5A). That is, the means of each condition fall within the 95% C. I. of the other conditions. Therefore, a fixational bias due to attentional cueing is not a sufficient explanation of our data.

**Figure 5.**
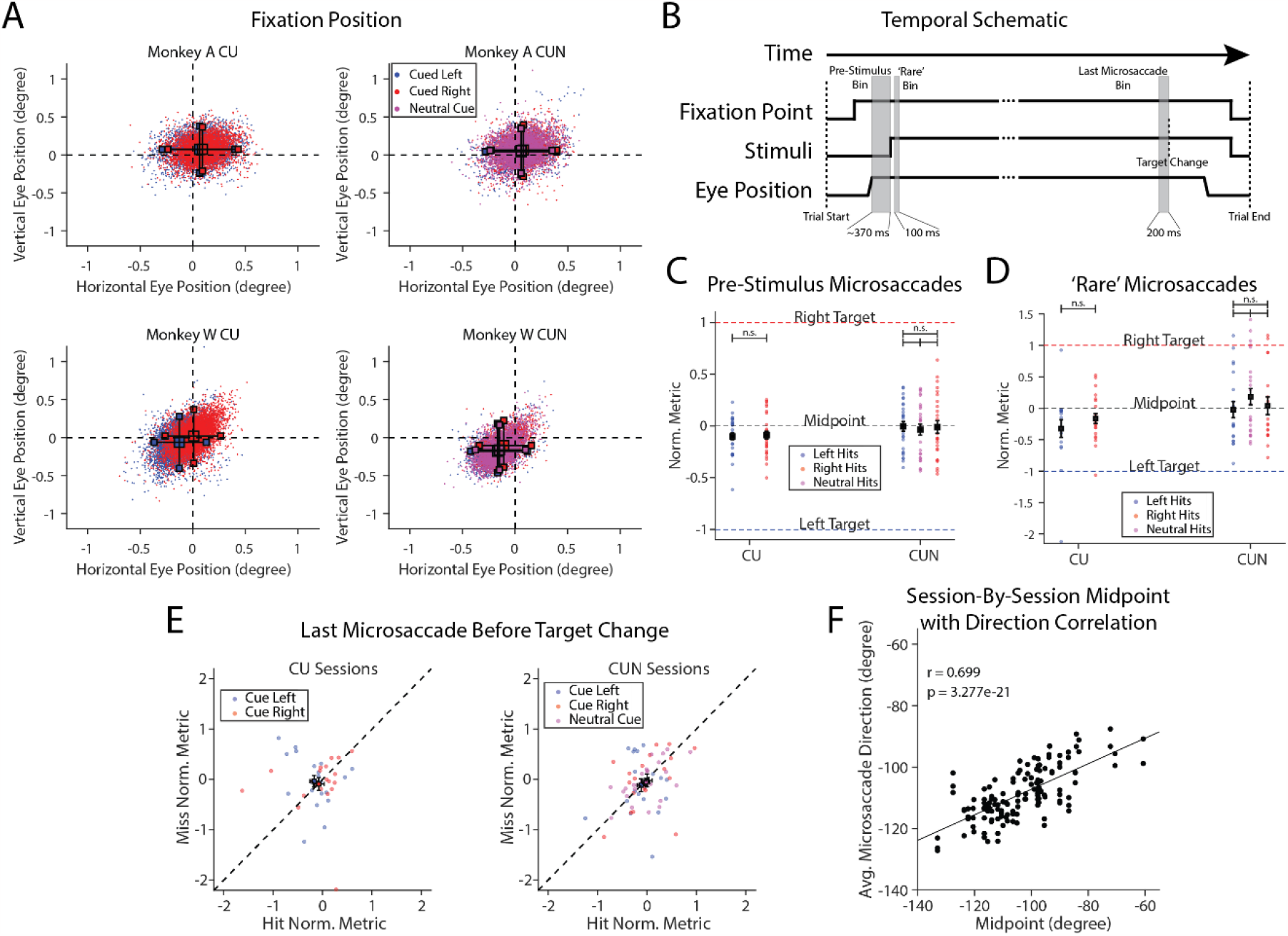
Alternative explanations fail to account for our results. A) Average fixation position by cueing regime (CU, left column; CUN, right column) and cued directions for monkey A (top row) and monkey W (bottom row). Individual points are the average fixation points per trial, large colored squares are the mean of the distribution and smaller squares connected by black lines indicate the 95% confidence intervals. The mean values of each distribution fall within the confidence intervals for the other conditions’ distributions across session type and monkey. B) A temporal schematic showing an example trial from start to finish. Gray shaded areas indicate different microsaccade analysis windows. Numbers below the bins indicate the bin duration. C) Normalization metric values for microsaccades that occur after monkeys acquire fixation but prior to stimulus onset (∼370-0 ms before stimulus onset). Same format as Figure 4B. Microsaccades are directed toward the midpoint and there are no significant differences in microsaccade direction between cued directions for CU or CUN cueing regimes. D) Normalization metric values for microsaccades that occur during peak microsaccade rate suppression (50-150 ms after stimulus onset). Same format as Figure 4B. Microsaccades are directed toward the midpoint, and there are no significant differences in direction between cued directions for CU or CUN cueing regimes. E) Normalization metric values for the single microsaccades that occur prior to target change (within -200 ms w.r.t target change), do not differentiate between hit and miss trials. Same format as Figure 4C. F) The session-by-session correlation between midpoint and microsaccade direction. Each point corresponds to a cueing condition from either CU or CUN regimes from either monkey (n = 136). r corresponds to the Pearson’s correlation and the solid line is the linear fit to the data. Note that the scale is larger in panels C & D than Figure 4B & C, respectively, likely because fewer microsaccades contribute to each data point.

Third, we investigated whether microsaccades were directed toward the midpoint between the two target positions could be caused by a low-level interaction of visual-motor representations in the superior colliculus (SC; 28). If the presence of visual stimuli activates two corresponding retinotopic locations in the SC, and those activated regions are subsequently “read out” or bias the impending motor command for the microsaccade, then we might expect microsaccades to be directed toward the midpoint of the two locations. To determine if such an interaction accounts for our results, we examined the direction of microsaccades after monkeys acquired fixation but prior to the presentation of the two stimuli (Fig. 1A, top panel & Fig. 5B). Using the same approach as in Fig. 4B, we found that the distribution of pre-stimulus microsaccades was centered around normalization metric values of 0, the midpoint (Fig. 5C). There were no significant differences between microsaccade direction across the CU or the CUN cueing regimes (CU Cued Left mean = -0.10 ± SEM = 0.03 vs. CU Cued Right mean = -0.09 ± SEM = 0.04; *p* = 0.62; CUN Cued Left mean = -0.01 ± SEM = 0.05 vs. CUN Cued Right mean = -0.01 ± SEM = 0.06; *p* = 0.89; CUN Cued Left vs. CUN Neutral Cue mean = -0.04 ± 0.05; *p* = 0.30; CUN Cued Right vs. CUN Neutral Cue; *p* = 0.55; ; Dunn-Bonferroni multiple comparison threshold, p = 0.0125). These data indicate that microsaccades were directed toward the target midpoint prior to visual stimuli onset, likely due to the maintenance of the cued location and target positions in memory.

Fourth, we investigated whether “rare but precious” (22) microsaccades during the microsaccade rate suppression (Fig. 2C, F) were more biased toward the cued target. Past work (22) has indicated that microsaccades that occur during this time window may be more informative about attentional state than other microsaccades. Therefore, we investigated the average microsaccade direction per session for microsaccades that occurred between 50-150 ms after stimulus onset (Fig. 5B). We removed conditions that had one or fewer microsaccades (total included session-condition combinations = 101/136). Using a similar approach as in Fig. 4B, we found that the distributions of microsaccade direction during the stimulus-related suppression were also centered around the normalization metric values of 0, the midpoint (Fig. 5D). There were no significant differences between microsaccade directions across the CU or the CUN cueing regimes (CU Cued Left mean = -0.32 ± SEM = 0.14 vs. CU Cued Right mean = -0.16 ± 0.08; *p* = 0.31; CUN Cued Left mean = -0.02 ± SEM = 0.13 vs. CUN Cued Right mean = 0.04 ± SEM = 0.14; *p* = 0.75; CUN Cued Left vs. CUN Neutral Cue mean = 0.18 ± SEM = 0.13; *p* = 0.27; CUN Cued Right vs. CUN Neutral Cue; *p* = 0.46; ; Dunn-Bonferroni multiple comparison threshold, p = 0.0125). These data indicate that microsaccades were still directed toward the midpoint in the time bin most associated with attentional biasing of microsaccades. Note that we failed to replicate the bias toward cued location in the CU data (Fig. 4B), likely because of the small number of microsaccades that occurred in this more limited time window.

Fifth, we asked if the relationship between microsaccade direction and the presumed locus of attention was obscured by overly large time windows of analysis. In theory, the microsaccade that occurs immediately before the stimulus change should be most closely associated with the locus of attention (17). We looked at the distribution of the last microsaccade prior to target change (within 200 ms) across hit and miss trials. If microsaccade direction reliably indicates the focus of attention, then we should expect to see microsaccades biased *towards* the detected target on hit trials and microsaccades biased *away* from the target on miss trials. However, we found that the distributions of the last microsaccade directions prior to target change did not significantly differ between hit and miss trials (Fig. 5E; CU Left Hit mean = -0.17 ± SEM = 0.08 vs CU Left Miss mean = -0.04 ± SEM = 0.12; *p* = 0.41; CU Right Hit mean = -0.08 ± SEM = 0.11 vs CU Right Miss mean = -0.09 ± SEM = 0.13; *p* = 0.94; CUN Left Hit mean = -0.12 ± SEM = 0.08 vs. CUN Left Miss mean = -0.11 ± SEM = 0.13; *p* = 0.98; CUN Right Hit mean = 0.00 ± SEM= 0.11 vs. CUN Right Miss mean = -0.02 ± SEM = 0.13; *p* = 0.88; CUN Neutral Hit mean = -0.02 ± SEM = 0.08 vs. CUN Neutral Miss mean = -0.05 ± SEM = 0.07; *p* = 0.63; Dunn-Bonferroni multiple comparison threshold, p = 0.01). These results indicate that, even when restricted to the window just prior to stimulus change, microsaccade direction was not different between hit and miss trials and therefore was unable to reveal the putative locus of covert attention.

Lastly, we investigated whether the observed midpoint bias was the result of a training induced bias. We reasoned that repeatedly performing the task might direct microsaccades towards the across-session average target position. To test this possibility, we correlated each session’s target midpoint with the corresponding average microsaccade direction for that session. If the midpoint bias is arbitrarily the result of training over many sessions, then the session midpoints and session microsaccade directions should be uncorrelated. On the other hand, if the microsaccade direction is sensitive to the specific target positions in each session, then the session midpoints and session microsaccade direction should be positively correlated. We found that the sessions’ average microsaccade directions were strongly correlated with the sessions’ midpoints (Fig. 5F, r = 0.699; p = 3.28 × 10^−21^), suggesting that our data are not the result of a training induced bias.

## Discussion

We investigated how variable cueing in a visuospatial attention task affected microsaccade direction. Across tens of thousands of microsaccades, the average microsaccade vector was not directed toward the cued target (Fig. 3), but instead toward the midpoint of the two potential targets (Fig. 4B). Microsaccades were slightly biased (∼10%) toward the cued location when using the more conventional CU cueing regime, consistent with the bias previously observed when an attentional cue is stored in memory (23). However, the bias dissipated when using the CUN cueing regime, suggesting that the addition of more ambiguously Cued blocks of trials reduced attentional modulation of microsaccade direction. Importantly, this attenuation in direction bias occurred across cueing regimes on trials with identical cue validity (i.e., the 80/20 Cued and Uncued trials). To further test the idea that microsaccade direction reveals the locus of covert attention, we compared the directions of microsaccades on hit versus miss trials. We found no difference in the direction of microsaccades between hit and miss trials (Fig. 4C). If attention was directed towards the midpoint, one would expect symmetrical performance on Cued and Uncued trials. However, this was not the case. Instead, we found canonical behavioral and neuronal effects of visuospatial attention, where performance was significantly better in Cued vs. Uncued trials (Fig. 1C & Mayo & Maunsell 2016). These results indicate that variable cue validity and implicit cues stored in memory weaken the microsaccade-attention relationship.

Why are microsaccades directed toward the midpoint of the visual stimuli? One potential explanation is that the monkeys simply happen to have a behavioral bias aligned to the midpoint of the target distributions. This explanation is unlikely for five reasons. First, there is a large range of possible directions in which a bias can occur (range of 180 deg. in our analyses). The bias in microsaccade direction that we observed was directed towards a meaningful location given the task demands of attending, to varying degrees, to both target locations. Second the session-by-session microsaccade direction was strongly correlated with each session’s target midpoint (Fig. 5F), suggesting the midpoint bias is dependent on session target position. Third, both animals showed a bias in the same direction despite no constraints beyond maintaining fixation for successfully performing the task. Fourth, the bias was more aligned to the midpoint than other salient directions, such as downwards. Finally, this explanation fails to account for the reduction of the cueing bias between the CU and CUN cueing regimes. We would expect a task irrelevant bias to be immune to changes in cueing validity, by definition. In total, these observations make the likelihood of a trivial behavioral bias unlikely.

One could also potentially argue that, based on Figure 3A, monkeys were targeting their microsaccades largely *away* from the midpoint. While it is true that many microsaccades fell along both directions of the midpoint axis (i.e., towards and away from the midpoint), it is difficult to conceive of a putative mechanism that improves visual acuity and cognitive enhancement in one visual hemifield by directing an eye movement to the opposite hemifield. It is more likely that compensatory eye movements were asymmetric because of idiosyncratic differences in square wave jerks, drift detection, and measurement noise. For example, monkey W appears to make more upward movements but also appears to have a larger downward fixation bias (Fig. 5A). Strictly speaking, we cannot rule out this possibility. Nevertheless, our central finding remains that microsaccades in our task fell along an axis that was not aligned with the targets and was well aligned to each session’s midpoint axis.

We considered several other possible explanations for midpoint-aligned microsaccades. Monkeys may show no meaningful cue-dependent differences in microsaccade direction because they shifted their fixation position dependent on cued direction. To see if this was the case, we measured the fixation position during the same time window that we measured microsaccades. We found no differences in the average fixation position across cued direction or session type (Fig. 5A). Therefore, drift in fixation position does not account for midpoint directed movements. Alternatively, competing neuronal representations of each target in brain regions underlying microsaccade generation (32) or the appearance of the targets themselves (2) may result in microsaccades directed toward the midpoint of two targets (28). To address these possibilities, we measured microsaccade directions prior to target onset. We found microsaccades were still directed towards the midpoint in the absence of visual stimuli (Fig. 5C), presumably due to the maintenance of the cued location and target positions in memory.

The fixation duration in the task was substantial (500-5000 ms), and we looked at the average direction across microsaccades in this epoch. It is conceivable that microsaccades appear to target the midpoint because we focused on this relatively large time window. For instance, the ‘rare but precious’ microsaccades that occur during the peak of the microsaccade rate suppression (22) are thought to show a larger bias towards the cued location. However, we found that even in this narrower time bin microsaccades were still directed toward the midpoint (Fig. 5D). Even the last microsaccade prior to target change did not differ in direction between hit and miss trials (Fig. 5E), indicating that in more constrained windows microsaccade direction remained insensitive to differences in the locus of attention.

An interesting possibility is that microsaccades along the midpoint minimize the eccentricity of potential targets. Therefore, movement along the midpoint axis in this task is a way in which subjects could hedge their bets and bring targets onto areas of the retina with higher acuity. If the goal of generating a microsaccade was to reduce behaviorally relevant target eccentricity, one may expect that as cue validity approaches 100%, microsaccade direction would align with the angle of the cued location. Indeed, we observed a bias of microsaccades toward the cued location with the more consistent cueing regime (CU) and an attenuation of bias with the less consistent cueing regime (CUN). The reduction in target eccentricity is seemingly the most parsimonious explanation of ours and previously reported results, and future work will need to test this hypothesis.

Seminal work by Engbert and Kliegl (2) used a cueing regime similar to our CUN cueing regime but found that, unlike our results, subjects’ microsaccade directions were biased toward the cued location. Why did they find a cue-related bias in their microsaccade directions, but we did not? A likely explanation points towards the differences in their task demands and stimulus locations. Engbert and Kliegl (2) presented a visual cue on every trial which may strengthen the information about cued location available. Indeed, recent work shows that explicit cues more strongly modulate microsaccade direction than implicit cues (23). Additionally, the task used by Engbert and Kliegl (2) presented the target at 0° or 180°. The alignment of the potential targets on the same axis is problematic for a number of reasons. First, the bimodal nature of microsaccades makes it difficult to determine if microsaccades towards fixation are directed at the fixation location or to the other potential target position. Our experimental design allows us to disambiguate these two options. Additionally in Engbert and Kliegl (2), the midpoint axis (the vertical meridian, in this case), is maximally distant to the stimulus axis, and the angular distance between these two axes may affect the production of microsaccades along that direction. The larger difference between the stimulus and midpoint axis may make it less likely that microsaccades will be generated along the midpoint. Future work will need to determine the interaction of stimulus position and attentional cueing in the modulation of microsaccade direction.

There is substantial literature suggesting that microsaccade direction reveals the locus of covert attention (1, 2, 17-19, 22). However, other groups (21, 33) have failed to find evidence to support this hypothesis, consistent with our results. What could account for the differences in results? One potential explanation is the difference in species. Our study, as well as that of Yu and colleagues (21), used non-human primates and did not find strong cue-dependent biasing of microsaccade direction. However, other groups have used non-human primates and found strong biasing of microsaccade direction (21), making the species difference an unlikely explanation. Continued work capitalizing on the benefits of both non-human primate and human subjects will be needed to pinpoint the factors that lead to microsaccades directed toward the target midpoint.

A more plausible explanation for the variability of results is differences in tasks and analyses. Experiments (1, 20) using 100% valid cues found strong biasing of microsaccade direction, which we failed to replicate with a more ambiguous cueing regime. Another key difference is that our task involved a cue stored in memory for a block of trials while other studies presented an explicit visual cue on each trail (1, 2, 20, 21). Furthermore, we controlled for anticipation by using fixation times sampled from a truncated exponential distribution (34), limiting temporal attention confounds. Finally, we analyzed the average angular direction of microsaccades and related that to target positions across cueing regimes, unlike studies (1, 2, 21) that quantified bias by comparing the distribution of microsaccade directions across cueing regime (e.g., attend in vs. attend out), with no relation to target position.

We chose our task parameters because in the real world it is unlikely for cues to be unambiguously predictive before salient events and to be predictable in time. Our results failed to support the hypothesis that microsaccade direction reveals the locus of covert attention, suggesting that the applicability of this idea to more naturalistic scenarios warrants additional research. Our results suggest that relatively straightforward task manipulations, such as the introduction of ambiguous cueing, can render the relationship between microsaccade direction and visual spatial attention tenuous. Indeed, the correlation across hit and miss directions (Fig. 4C) implies that the neuronal populations responsible for generating microsaccades may be robust to attentional effects. Future work should probe the attentional modulation of firing rates in the SC, a key site for the generation of microsaccades (32).

Considering the literature on microsaccades more broadly, it is perhaps not surprising that microsaccade directions did not “point” to the attended targets in our tasks. In some ways, microsaccade direction appears to be a particularly frail measure of attention. Microsaccades do not occur in a significant portion of trials in cued attention tasks. For example, in Hafed and Clark (1) human subjects did not make microsaccades in roughly 75% of cue periods. In our dataset monkeys did not make microsaccades in nearly 40% of trials. Microsaccade direction has also been shown to depend on stimulus changes (2), stimulus type/stimulus arrangement (28), and may depend on biases in fixation position (31). Even one of the more convincing and clever experiments done to probe the relationship between microsaccades and attention, in which subject performance was measured time-locked to spontaneous microsaccades, did not show consistent benefits of microsaccades for every subject (17). Finally, it is important to note that the majority of studies, including our own, use impoverished foveal stimuli (e.g., a fixation point) (35). During natural viewing conditions, more complex fixational stimuli are more likely to elicit exploratory microsaccades (36), likely further increasing the independence between microsaccade direction and covert attention. Taken together, these factors complicate the interpretation that microsaccade direction can be used as a measure of latent neuronal states.

How do our results affect the usefulness of microsaccades as a biomarker? Although our results call into question the use of microsaccade *direction* as a generalizable, overt measure of covert attention, there are other features of microsaccades that may be used as biomarkers for cognitive states or neurological disease. For example, microsaccade *amplitude* may relate to covert attention (37). Additionally, microsaccade *rate* may prove a more useful parameter to reveal cognitive fluctuations in internal state (38). Although we found no clear indication of this in our data (not shown), future work is needed to determine if these other parameters, individually or in combination, reveal the locus of covert attention in more dynamic scenarios.

Our results suggest that microsaccade direction may not be useful as a biomarker for attentional state. But microsaccades may be valuable in diagnosing neurological disorders (16). These disorders are often sensitive to disruptions in oculomotor pathways and therefore may alter microsaccade dynamics. An interesting possibility is that microsaccades are useful as biomarkers because they modulate ongoing neuronal processing. For example, recent work found that microsaccades affect the attentional modulation of cortical (20, 39) and subcortical activity (21). Additionally, microsaccades modulate primary visual cortical activity and ongoing visual experience in a direction specific way (40). It is imperative that future work disentangles the mechanisms by which microsaccades subserve diagnosis of neurological disease. Not only could it improve and constrain the interpretations of current microsaccade-related biomarkers, but it could also facilitate development of new ways in which microsaccades could reveal covert neuronal states.

## Materials and Methods

Surgical details and task information are detailed in our prior work (3). This work contains a superset of those data using the same methods, and details relevant to the current investigation are below.

### Subjects

Two male *Macaca mulatta* monkeys were surgically implanted with titanium headposts and scleral eye coils for the tracking of eye movements (24). Surgeries were performed under isoflurane anesthesia with standard courses of analgesics, antibiotics, and anti-inflammatories and with approval of the Institutional Animal Care and Use Committee of Harvard Medical School.

### Behavioral Task

Monkeys were trained to detect an orientation change in one of two simultaneously presented stimuli. The task began when the monkey fixated a small white square at the center of the CRT monitor. Two Gabor stimuli then appeared and remained on the screen until the end of the trial. After a random amount of time (500 – 5500 ms, mean 3000 ms, truncated exponential distribution), one of the stimuli changed in orientation (0.3°-60°). The monkey was rewarded for reporting the change by making a saccade to the Gabor within 100-550 ms after the orientation change. Only one target could change on any given trial, and the monkey was rewarded for reporting the change regardless of the location of the change. Five percent of trials were catch trials where neither target changed, and the monkey was rewarded for maintaining fixation until the end of the trial. To facilitate neuronal recordings in visual area V4 (not analyzed here; see 3), stimuli were presented in the lower half of each hemifield.

We cued attention in two different regimes: the Cued/Uncued (CU) regime and the Cued/Uncued/Neutral (CUN) regime. The CU regime consisted of ∼50 trials per block in which a single location was cued in each block. The target change was 80% likely to occur at the cued location (“Cued” trials) while the target change was 20% likely to occur at the uncued location (“Uncued” trials), hence the name Cued/Uncued. The CUN regime consisted of ∼50 trials per block in which either one target location was cued or other blocks in which both target locations were cued (3). When a single location was cued, the target change was 80% likely to occur at the cued location while the target change was 20% likely to occur at the uncued location. When both target locations were cued (‘Neutral” trials), the target change was 50% likely to occur at either target location, though on any given trial only a single target would change. Therefore, the CU regime consisted of blocks with 80% cue validity and the CUN regime consisted of blocks with 80/50% cue validities. Only a single cueing regime was used during a given recording session. Trial locations were explicitly cued with brief white flashes over the target(s) at the start of each block of trials (“instruction trials”). On instruction trials, the cue appeared over the target that would change 100% of the time in the CU regime and would appear over both targets in the CUN regime. Monkeys were required to complete ∼5 instruction trials (data not analyzed) before proceeding with the analyzed block of trials with no explicit cueing (3).

We analyzed the behavioral performance and eye movements from 29 recording sessions (14 for monkey A, 15 for monkey W) that alternated between Cued blocks to the left and right stimulus locations (CU cueing regime). To probe the effect of more variable cueing, we analyzed an additional 26 sessions (14 for monkey A, 12 for monkey W) that consisted of equal proportions of Cued and Neutral blocks of trials (CUN cueing regime), which alternated Right-Neutral-Left-Neutral-Right and so on.

Proportion correct corresponds to the ratio of hits versus misses for a given condition:

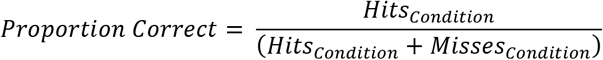

Where ‘condition’ corresponds to the particular cueing condition (Cued, Uncued, or Neutral) and target location (left or right). The behavioral performance of the monkeys on CUN sessions has been reported previously (3). Behavioral performance scales as a function of cueing condition, in agreement with human psychophysical results (41-46).

### Stimuli

Odd-symmetric, full-contrast Gabor stimuli were sinusoidally counterphased at 10 Hz. Gabors ranged in size, defined by the standard deviation to reduce edge effects, from σ = 0.45° -1.6°. Gabors ranged in spatial frequency (0.3-2.2 cycles/deg) and initial orientation determined by the preferences of a simultaneously recorded neuronal population (not discussed in this manuscript). Stimuli were presented on a CRT monitor (Frame rate = 100 Hz, pixels = 1024 × 768, bits/pixel = 32, 8-bit DACs) and displayed on a uniform gray background. This monitor was calibrated to produce linear steps in luminance and positioned 57 cm from the subject.

Target positions were chosen to align with the neuronal receptive fields of implanted electrode arrays. Leftward targets were presented from ⊖ ≈ -164° to -108° and rho = 5.22° to 27° for monkey A and ⊖ ≈ -171° to -132° and rho = 4.61° to 8.32° for monkey W (Fig. 1B). Rightward targets were presented from ⊖ ≈ -77.9° to -33° and rho = 4.47° to 18.1° for monkey A and ⊖ ≈ -71.6° to -20.2° and rho = 6.27° to 10.3° for monkey W (Fig. 1B). Therefore, left target eccentricity ranged from 4.61° to 27° and right target eccentricity ranged from 4.47° to 10.3°.

### Detection of Eye Movements

Eye position was sampled at 200 Hz via a scleral eye coil. Eye position signals were converted to polar coordinates, and radial velocity was calculated by differentiating the radial magnitude time series for each trial (47). The radial velocity time series was smoothed using a 15 ms (3 sample) rectangular filter and the absolute value of the time series were taken. Samples of the radial velocity time series were classified as belonging to an eye movement if at least two consecutive samples exceeded the velocity threshold of 6° per second. Eye movement onset was defined as the first sample when eye velocity increased above the sum of the mean and standard deviation of the radial velocity time series (evaluated with eye movement samples removed). Eye movement offset was defined as the second consecutive sample with a velocity lower than the sum of the mean and standard deviation of the radial velocity time series (evaluated with eye movement samples removed).

To prevent potential experimenter bias introduced by manual curation and to facilitate replicability across studies, we automatically removed spurious eye movements by implementing an exclusion metric. The metric was based on the natural logarithm of the ratio between the total displacement of the movement and the path length of the movement:

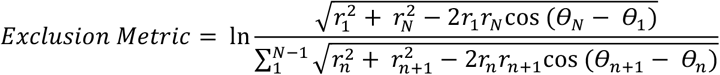

Where N corresponds to the number of samples in the detected eye movement, n corresponds to the sample position in the N time series, r corresponds to the rho (radial magnitude), and ⊖ corresponds to the angular position. The validity of the exclusion metric rests on the assumption that saccadic eye movements are relatively straight. Straight movements would result in a value of zero because the path length and total displacement are equivalent. Movements were excluded if this metric exceeded one (2.4% of movements).

Detected eye movements that passed all criteria were then labeled as microsaccades if the total displacement of the movement was less than 2° and labeled as saccades if the displacement was greater than or equal to 2°. We saw the same effects when using a 1° or 0.5° cutoff. All analyses were done on microsaccades during fixation, after the stimulus onset and prior to the target change (gray shading in Fig. 1C) unless otherwise noted. We only analyzed correctly detected (“Hit”) validly Cued trials (trials in which the cued target changed and the monkey made an eye movement to it) unless otherwise noted in the main text. In total we analyzed 22,097 microsaccades from 17,813 valid hit trials in monkey A and 72,050 microsaccades from 21,355 valid hit trials in monkey W.

### Analysis of Microsaccades

#### Microsaccade Distributions

The goal of the analyses was to quantify the microsaccade direction distributions. Therefore, each microsaccade was centered so that the starting position of the movement was the origin in polar coordinates. The angular endpoints of microsaccades for a given session, or condition, were binned in 10° bins between [-180°:10°:180°]. To facilitate statistical inference, we reduced the modality of these distributions by selecting the microsaccades that moved the fovea into the hemifield containing both targets. We did this by dividing the visual field along the axis orthogonal to the Euclidian midpoint of the two targets. Circular mean and resultant vector length were computed through MATLAB’s Circular Statistics Toolbox (30). Unless otherwise noted, analyses were performed on these selected distributions.

#### Normalization Metric

To quantify the directional bias across sessions that contained slightly different target positions, we normalized the circular mean of the microsaccade distribution to the axis connecting the two targets:

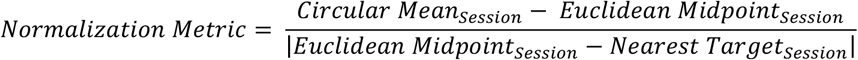

Where the nearest target corresponds to the target (left vs. right) closest to the circular mean of the session. That is, if the microsaccade was left of the midpoint the microsaccade direction was normalized by angular difference between the left target and the midpoint (⊖_L_, Fig. 4A) or was normalized by the angular difference between the right target and midpoint (⊖_L_, Fig. 4A) if the microsaccade was to the right of the midpoint. Therefore, values of -1 correspond to a circular mean at the same angular position of the left target, values of 0 correspond to a circular mean that aligns with the Euclidean midpoint between the two targets, and values of 1 correspond a circular mean at the same angular position of the right target. We combined data across monkeys for the normalization metric due to the similarity in their behavior. The normalization metric only standardizes the direction of microsaccades across monkeys and target positions but does not account for differences in microsaccade number across monkeys.

#### Statistical Analyses

We performed a series of two-tailed paired t-tests to compare the majority of the distributions. The exception was for Figure 5C in which unpaired t-tests were used due to the different number of sessions per condition. We accounted for multiple comparisons with a Dunn-Bonferroni correction. We correlated session-by-session midpoint with session-by-session microsaccade direction using MATLAB’s *corr* function and performed the linear fit to the data with the *fitlm* function.

## Acknowledgments

This work was supported by National Institutes of Health Grant R21EY033460, CORE Grant P30EY08098 to the Department of Ophthalmology, the Eye and Ear Foundation of Pittsburgh, and unrestricted funds from Research to Prevent Blindness, New York, NY. We thank John Maunsell for the use of data collected in his laboratory. We thank members of the Mayo and Gandhi laboratories, Christopher Henry, and Matthew Smith for feedback on this work.

